# Genetic and epigenetic determinants of reactivation of Mecp2 and the inactive X chromosome in neural stem cells

**DOI:** 10.1101/2021.02.25.432827

**Authors:** Hegias Mira-Bontenbal, Beatrice Tan, Cristina Gontan, Sander Goossens, R.G. Boers, J. Boers, Catherine Dupont, Martin van Royen, Wilfred van IJcken, Pim French, Toni Bedalov, Joost Gribnau

## Abstract

Rett Syndrome is a neurodevelopmental disorder in girls that is caused by heterozygous inactivation of the chromatin remodeler gene *MECP2*. Rett Syndrome may therefore be treated by reactivation of the wild type copy of *MECP2* from the inactive X chromosome. Most studies that model *Mecp2* reactivation have used mouse fibroblasts rather than neural cells, which would be critical for phenotypic reversal, and rely on fluorescent reporters that lack adequate sensitivity. Here, we present a mouse model system for monitoring Mecp2 reactivation that is more sensitive and versatile than any bioluminescent and fluorescent system currently available. The model consists of neural stem cells derived from female mice with a dual reporter system where MECP2 is fused to NanoLuciferase and TdTomato on the inactive X chromosome. We show by bioluminescence and fluorescence that *Mecp2* is synergistically reactivated by 5-Aza treatment and *Xist* knockdown. As expected, other genes on the inactive X chromosome are also reactivated, the majority of which overlaps with genes reactivated early during reprogramming of mouse embryonic fibroblasts to iPSCs. Genetic and epigenetic features such as CpG density, SINE elements, distance to escapees and CTCF binding are consistent indicators of reactivation, whereas different higher order chromatin areas are either particularly prone or resistant to reactivation. Our MeCP2 reactivation monitoring system thereby suggests that genetic and epigenetic features on the inactive X chromosome affect reactivation of its genes, irrespective of cell type or procedure of reactivation.

## Introduction

Rett syndrome (RTT) is the second most prevalent cause of intellectual disability in girls after Down Syndrome, affecting 1 in 10,000 live female births (1). It is caused by heterozygous mutations in the methyl-CpG-binding protein 2 (MECP2), whose gene is X-linked and subject to random X chromosome inactivation (XCI) during early embryogenesis. RTT-affected girls are thus mosaic in terms of *MECP2* expression: half of their cells will express the wild type (WT) copy of *MECP2*, while the other half will express the mutant *MECP2* allele. This also implies that RTT-affected cells have a silenced WT *MECP2* copy located on the inactivated X chromosome (Xi). Previous work has shown that postnatal re-expression of WT *Mecp2* copies in a RTT mouse model reverts its phenotype (2, 3), which has led major interest in the RTT field to re-express WT MECP2 in human RTT patients. One way of achieving this is by reactivation of the endogenous WT copy of *MECP2* on the Xi in RTT cells. To revert inactivation of an X-linked gene, one needs to understand the different steps that happen during XCI. Cells in the inner cell mass of the female mouse blastocyst bear two active X chromosomes. Upon differentiation and epiblast formation, one of the X chromosomes is randomly chosen to upregulate expression of the long non-coding RNA *Xist*. This results in the coating of a single X chromosome with *Xist* and the recruitment of proteins such as SPEN, RBM15, HDAC3, and the polycomb repressive complexes PRC1 and PRC2 to silence X-linked genes in cis. Eventually, CpGs at promoters become methylated to lock XCI down.

Several studies have delved into the mechanics of *Mecp2* reactivation or in more general terms, X chromosome reactivation (XCR) in mouse cells and tissues by either looking for factors that are important to maintain *Xist* expression, by directly knocking down *Xist* or by inhibiting the DNA methyltransferase DNMT1 (4–9). The combination of *Xist* knockdown using shRNA or antisense oligonucleotides (ASOs) with 5-Azacitidine (5-Aza, a DNMT1 inhibitor) treatment synergistically reactivated *Mecp2* fused to a firefly luciferase reporter on the Xi of a mouse fibroblast cell line (5, 8). In addition, blocking the PI3K/AKT/mTor pathway using an inhibitor of SGK1, a downstream effector of PDPK1, or mTOR with GSK650394 or rapamycin respectively, results in biallelic expression of *Mecp2* in mouse fibroblasts, while inhibition of ACVR1 with LDN193189 leads to similar results (7). Treatment of different fibroblasts carrying a GFP transgene on the Xi with rapamycin, GSK650394 or LDN193189 led to increased fluorescence (7), confirming that the PI3K/AKT/mTOR and BMP pathways are involved in maintenance of repression of the Xi. *In vivo*, injection of GSK650394 and LDN193189 in brains of *Xist ^-/+^:Mecp2^+/GFP^* mice where *Mecp2* is fused to GFP on the Xi, resulted as well in significant GFP expression (7). Also, inhibition of DNMT1 and Aurora kinases led to synergistic reactivation of an Xi-linked GFP transgene (9). A more suitable approach to perform high throughput chemical compound screens for *Mecp2* reactivation requires the generation of an improved mouse model and the derivation of its associated cell lines closer to the neuronal target cells and the usage of a highly sensitive luciferase, instead of fluorescence, whose expression is under the control of the endogenous *Mecp2* promoter and not a transgene on the X chromosome.

Here, we have developed a mouse model system where *Mecp2* is fused to NanoLuciferase, a luciferase enzyme smaller and 100 times brighter than the regular firefly luciferase. We have also introduced a fluorescent TdTomato reporter downstream of NanoLuciferase separated by a P2A signal. This dual capability not only permits measurement of NanoLuciferase activity at a populational level, but also measurement of Tomato fluorescence at the single-cell level. Our *Xist^-/+^:Mecp2^+/NLucTom^* compound mice display complete skewed XCI of the reporter allele and are generated in a highly polymorphic C57BL/6:Cast/Eij (maternal:paternal) F1 hybrid background providing a wealth of SNPs for X-chromosome-wide allele-specific expression analysis. From these mutant mice, we have isolated mouse embryonic fibroblasts (MEFs), embryonic stem cells (ESCs) and neural stem cells (NSCs) for further studies and to provide them to the community. We show that 5-Aza treatment in combination with *Xist* knockdown in NSCs leads to XCR with striking resemblance to iPSC-reprogramming-specific XCR (10), suggesting a general pattern in the capability of X-linked genes to reactivate independently of the mechanism. In this manuscript, we highlight the potential of our model to study XCR.

## Results

### Generation of *Mecp2-NanoLuciferase-TdTomato* mice

To obtain highly polymorphic *Xist^-/+^:Mecp2^+/NLucTom^* mice, we firstly generated *Mecp2^NLucTom/Y^* ESCs in a Cast/EiJ (cast) background. We transfected WT male cast ESCs with the NanoLuciferase-P2A-TdTomato construct where NanoLuciferase is fused to the C-terminus of *Mecp2* and TdTomato (Tomato from hereon) is translated as an independent protein thanks to a P2A self-cleaving peptide (Fig. 1*A*). FACS analysis showed a distinct Tomato-positive cell population that was sorted and expanded (Fig. 1*B*). PCR analysis using primers spanning the 5’ and 3’ specific integration sites and primers against the endogenous allele confirmed proper integration on DNA obtained from sorted Tomato-positive cells (Fig. 1*C*, Table S1). This resulted in the appearance of a higher molecular weight band of MECP2 by immunoblotting due to its fusion to NanoLuciferase (19 kDa, Fig. 1*D*). Luminescence analysis showed very strong NanoLuciferase activity in *Mecp2^NLucTom/Y^* ESCs compared to WT ESCs (Fig. 1*E*). Cells were then injected in blastocysts and a cast colony of *Mepc2-NLucTom* mice was generated. *Mecp2^NLucTom/Y^* mice are viable with normal lifespan and do not show any RTT-related phenotype, indicating that the fusion of NanoLuciferase to MECP2 is not deleterious to its function (Fig. S1*A*). Immunofluorescence (IF) for NanoLuciferase and Tomato fluorescence analysis in *Mecp2^+/LucTom^* female brains show that MECP2-NanoLuciferase and Tomato are expressed in 50% of the cells as expected from random XCI (Fig. S1*B*). Moreover, *Mecp2^+/LucTom^* female brains also show high NanoLuciferase activity compared to WT controls, highlighting the usefulness of this system for *in vivo* studies (Fig. S1*C*). We have thus generated a *Mecp2*-NanoLuciferase-Tomato mouse colony in a cast background.

**Fig. 1.**
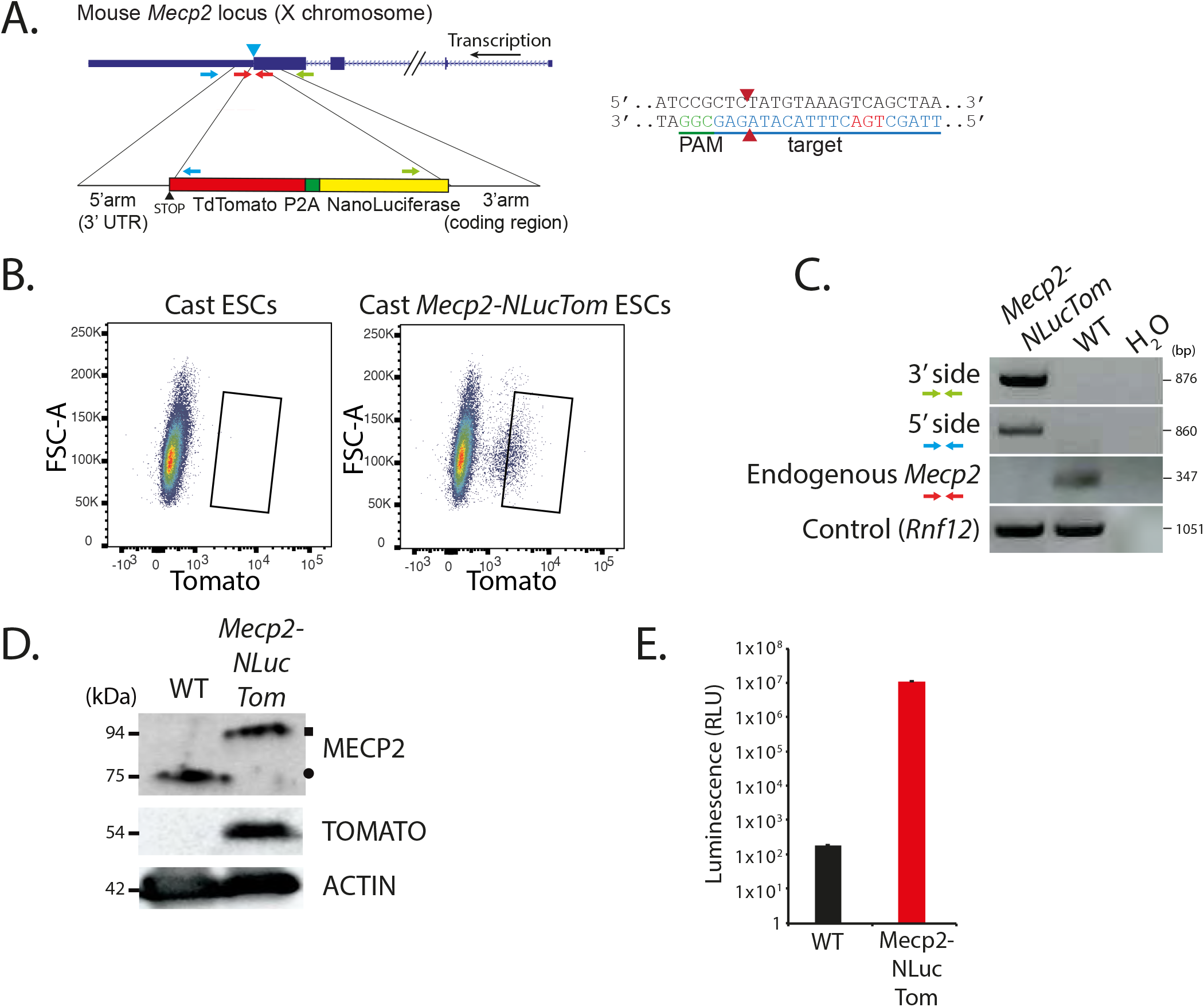
*Mecp2^NLucTom/Y^* male cast ESCs show proper reporter integration and expression. (***A***) Mouse *Mecp2* locus with the NanoLuciferase and Tomato donor vector. Green, blue and red primer sets were used to amplify the 3’ integration site (forward primer in NanoLuciferase, and reverse primer outside of 3’ homology arm), the 5’ integration site (forward primer outside the 5’ homology arm, and reverse primer inside Tomato) and non-targeted endogenous end of *Mecp2* respectively (see Fig. 1*C*). The guide RNA, PAM, cutting site (red arrows) and *Mecp2’s* TGA STOP codon are depicted on the right. Primer sequences are found in Table S1. (***B***) FACS plots depicting Tomato fluorescence before and after transfection of WT male cast ESCs with the CRISPR/Cas9 and donor vector depicted in (***A***). The rectangle shows the sorted population. (***C***) Genomic PCR with primers described in (***A***) on FACS-sorted *Mecp2^NLucTom/Y^* ESCs and parental WT ESCs showing proper integration of the reporters. A control locus PCR band is depicted *(Rnf12). (**D**)* Western Blot analysis of FACS-sorted *Mecp2^NLucTom/Y^* ESCs and parental WT ESCs showing a shift in the molecular weight of MECP2. Tomato is translated as an independent protein thanks to the P2A signal. Actin was used as loading control. MECP2-NLuc and WT MECP2 are indicated by a square and a circle, respectively. (***E***) NanoLuciferase activity assay showing the expression of NanoLuciferase from sorted *Mecp2^NLucTom/Y^* ESCs (500,000 cells analysed per well, average activity ± s.d., n=3 biological replicates).

### Generation of *Xist^-/+^:Mecp2^+/NLucTom^, Xist^-/+^:Mecp2^-/NLucTom^, Mecp2^+/NLucTom^* and *Mecp2^NLucTom/NLucTom^* cell lines

To study *Mecp2* reactivation, we crossed cast *Mecp2^NLucTom/Y^* males with C57BL/6 (Bl6) WT, *Zp3-Cre:Xist^2lox/2lox^* or *Zp3-Cre:Xist^2lox/2lox^:Mecp2^2lox/2lox^* females (Fig. S2*A*). Oocyte-specific expression of Cre, thanks to *the Zp3-Cre* transgene, results in recombination of loxP sites before fertilization, resulting in embryos that are *Xist^-/+^:Mecp2^+/NLucTom^* and *Xist^-/+^:MecpZ^-/NLucTom^* (among other genotypes). In this way, we isolated Bl6:cast WT controls, *Xist^-/+^:Mecp2^+/NLucTom^, Xist ^-/+^:Mecp2^-/NLucTom^, Mecp2^+/NLucTom^* and cast *Mecp2^NLucTom/NLucTom^* MEF, ESC and NSC lines. We confirmed by IF SOX2 expression and absence of the differentiated neuron-specific marker TUJ-1 in our NSC lines (Fig. S2*B*). Our NSC lines were also able to differentiate into TUJ-1-expressing neurons, GFAP-expressing astrocytes and OLIG2-expressing oligodendrocytes, confirming stemness (Fig. 2*A*). *In vitro* grown neurons with RTT (*Xist^-/+^:MecpZ^-/NLucTom^*) did not show any differences compared to WT or *Xist^-/+^:Mecp2^+/NLucTom^* neurons in terms of nuclear size, number of roots and extremities per nucleus and total neurite length per nucleus (Fig. S2*C*).

**Fig. 2.**
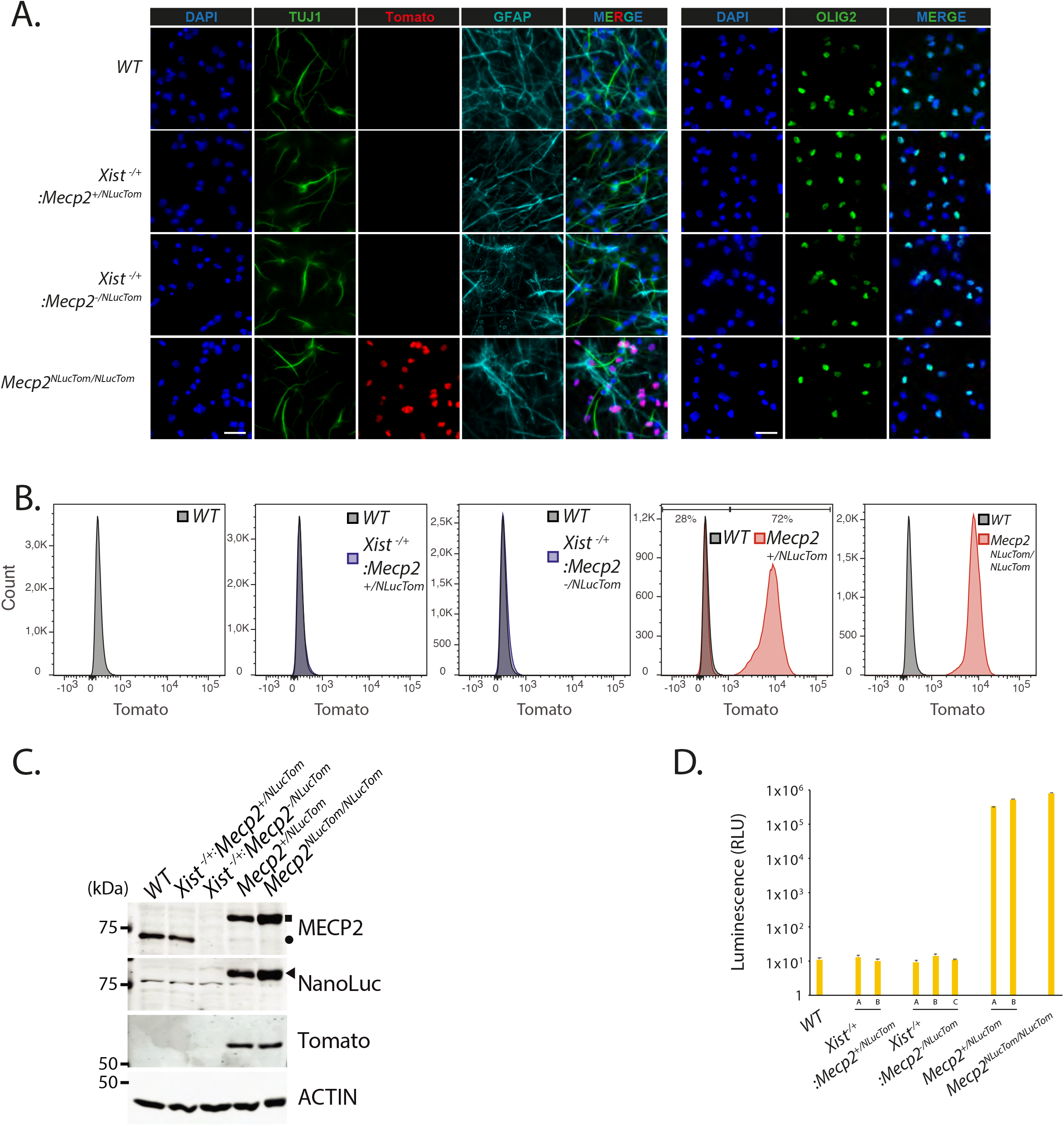
Characterization of *Xist^-/+^:Mecp2^+/NLucTom^, Xist^-/+^:Mecp2^-/NLucTom^, Mecp2^+/NLucTom^* and *Mecp2^NLucTom/NLucTom^* NSCs. (***A***) IF of TUJ1 (green left), GFAP (turquoise) and OLIG2 (green right) of WT, *Xist^-/+^:Mecp2^+/NLucTom^, Xist^-/+^:Mecp2-^MLucTom^* and *Mecp2^NLucTom/NLucTom^* NSCs differentiated towards neurons, astrocytes and oligodendrocytes. Tomato fluorescence was measured directly. Blue, DAPI. White scale bars: 25 μm. (***B***) FACS analysis of Tomato fluorescence of WT, *Xist^-/+^:Mecp2^+/NLucTom^, Xist^-/+^:Mecp2^-/NLucTom^* and *Mecp2^NLucTom/NLucTom^* NSCs. The percentage of Tomato-positive *Mecp2^+/NLucTom^* NSCs is shown. (***C***) Western Blot analysis of WT, *Xist ^-/+^:Mecp2^+/NLucTom^, Xist ^-/+^:Mecp2 ^NLucTom:^* and *Mecp2^NLucTom/NLucTom^* NSCs showing expression of MECP2-NLuc in *Mecp2^+/NLucTom^* and *Mecp2^NLucTom/NLucTom^* NSCs but not in *Xist^-/+^:Mecp2^+/NLucTom^, Xist^-/+^:Mecp2^-/NLucTom^* NSCs as expected. MECP2-NLuc, WT MECP2 and NanoLuciferase are indicated by a square, a circle and a triangle respectively. Tomato is expressed as an independent protein. (***D***) NanoLuciferase activity assay of several clones of WT, *Xist^-/+^:Mecp2^+/NLucTom^*, *Xist ^-/+^:Mecp2^-/NLucTom^, Mecp2^+/NLucTom^* and *Mecp2^NLucTom/NLucTom^* NSCs showing an increase of 4-to-5 levels of magnitude of NanoLuciferase activity from an active X chromosome in *Mecp2^+/NLucTom^* and *Mecp2^NLucTom/NLucTom^* NSCs (50.000 cells were analysed per clone per well, average activity ± s.d., n=3 biological replicates).

Full skewing of XCI of the paternal cast allele in *Xist^-/+^:Mecp2^+/NLucTom^* and *Xist^-/+^:Mecp2 ^-/NLucTom^* NSCs was confirmed by FACS analysis (Fig. 2*B*). In addition, *Mecp2^+/NLucTom^* NSCs displayed skewed XCI as expected from their hybrid origin, where around 60-70% of the cells show inactivation of the Bl6 allele (11). *Mecp2^NLucTom/NLucTom^* NSCs displayed a single Tomato-positive peak. Completely skewed XCI in *Xist^-/+^:Mecp2^+/NLucTom^* and *Xist^-/+^:MecpZ^-/NLucTom^* NSCs and absence of *Mecp2* expression in *Xist^-/+^:MecpZ^-/NLucTom^* NSCs was also demonstrated by immunoblotting analysis (Fig. 2*C*). *Mecp2^+/NLucTom^* and *Mecp2^NLucTom/NLucTom^* NSCs showed a higher molecular weight band for MECP2 protein, NanoLuciferase and Tomato expression. Finally, NanoLuciferase activity analysis showed that several *Mecp2^+/NLucTom^* and *Mecp2^NLucTom/NLucTom^* NSC clones have high levels of NanoLuciferase activity, and as expected, several *Xist^-/+^:Mecp2^+NLucTom^* and *Xist ^-/+^:Mecp2 ^NLucTom^* NSC clones do not (Fig. 2*D*). The background levels of NanoLuciferase expression in *Xist^-/+^:Mecp2^+/NLucTom^* and *Xist^-/+^:Mecp2 ^-/NLucTom^* NSCs are similar to WT cells indicating that escapism of *Mecp2-NanoLuciferase* from the inactive cast X chromosome is virtually inexistent. To quantify the level of transcriptional repression of *Mecp2-NLuc-Tom* on the inactive X-chromosome, we compared the NanoLuciferase activity in cells with the reporter on the active and inactive X. The reporter exhibited >30,000 lower activity when on the inactive X chromosome compared to the active X chromosome (Fig. 2*D*).

### Reactivation of the inactive *Mecp2*-NanoLuciferase allele

The compounds LDN193189 and GSK650394, which inhibit ACVR1 and SGK1 respectively, have been shown to reactivate an inactive GFP reporter on the Xi chromosome in fibroblasts and an inactive *Mecp2-GFP* fusion gene in mouse brains (7). In addition, the HDAC1/3 inhibitor RG2833 has been shown to facilitate XCR during reprogramming of female Xi-linked GFP transgenic MEFs (10).

We therefore treated our NSCs with LDN193189, GSK650394, RG2833 and/or decitabine (structurally very similar to 5-azacitidine, and called 5-Aza henceforth) for seven days. Single treatments with LDN193189, GSK650394 or RG2833 and double treatment with LDN193189 or GSK650394 did not result in *Mecp2* reactivation (Fig. 3*A*). Double treatment of LDN193189 or RG2833 with 5-Aza showed reactivation of the silent NanoLuciferase reporter comparable to single treatment with 5-Aza, indicating that 5-Aza is the only tested drug that reactivates the silent copy of *Mecp2*.

**Fig. 3.**
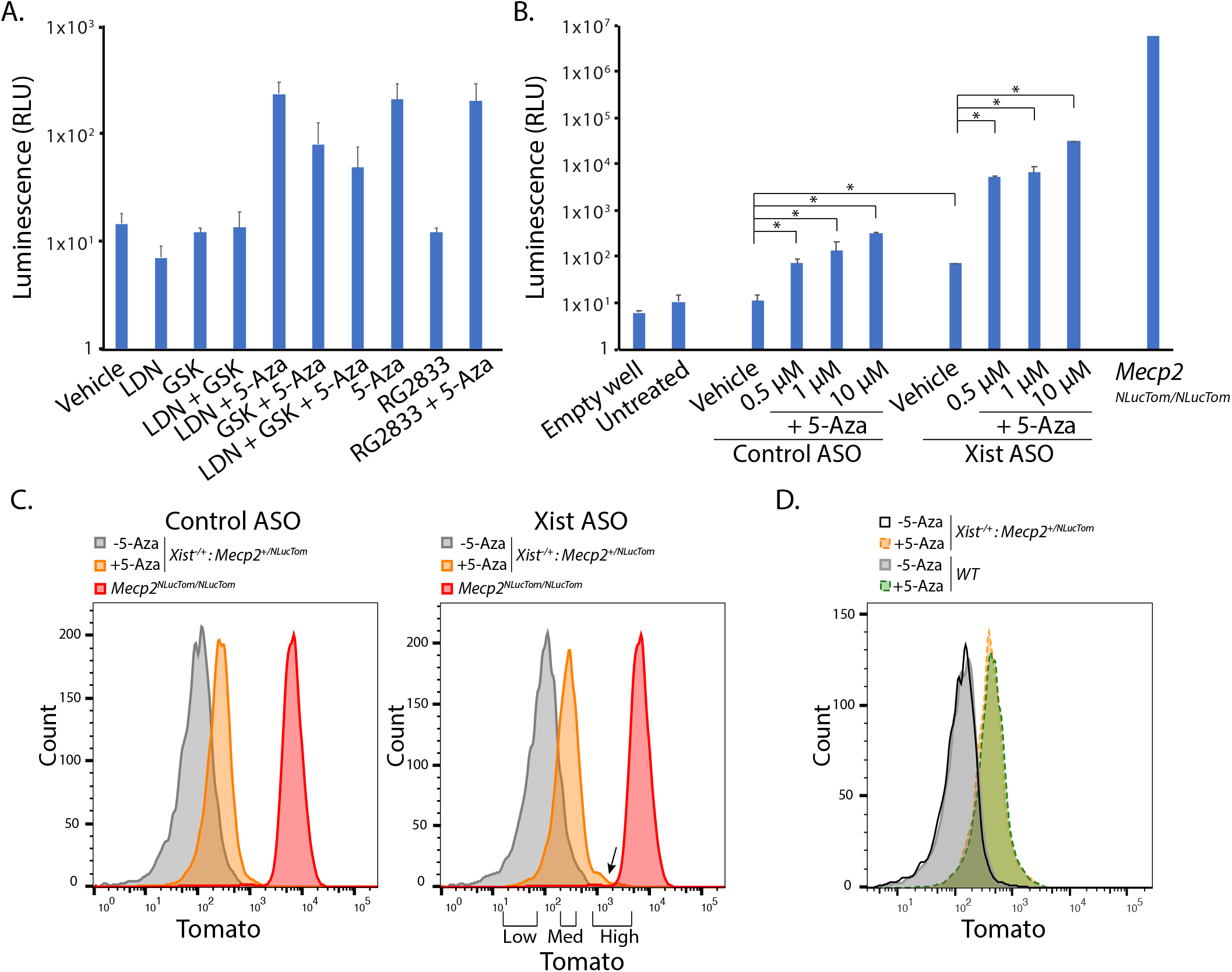
5-Aza treatment of *Xist^-/+^:Mecp2^+/NLucTom^* NSCs leads to reactivation of the NanoLuciferase-Tomato dual reporter. (***A***) NanoLuciferase activity assay of *Xist^-/+^Mecp2^+/NLucTom^* NSCs treated with LDN193189 (LDN), GSK650394 (GSK), RG2833 or 5-Aza in different combinations for 7 days (300,000 cells per well, average activity ± s.d., n=3 biological replicates). (***B***) NanoLuciferase activity assay of *Xist ^-/+^:Mecp2^+/NLucTom^* NSCs treated with different concentrations of 5-Aza in combination with control ASOs or *Xist* ASOs days (average activity ± s.d., n=3 biological replicates). Significant differences are depicted with an asterisk (500.000 cells were analysed per well, Two-tailed Student t-test, p-value < 0.05). (***C***) FACS plots depicting *Xist^-/+^:Mecp2^+/NLucTom^* NSCs treated with control or *Xist* ASOs with (orange) or without (gray) 5-Aza for 3 days. *Mecp2^NLucTom/NLucTom^* NSCs are shown as Tomato-positive controls (red). FACS-sorted populations that were subsequently analysed by RNA-se are shown (Low, Medium, High). The shoulder in Xist ASO and 5-Aza-treated sample is shown by an arrow, this corresponds to the Tomato-High population. (***D***) FACS plots depicting WT and *Xist^-/+^:Mecp2^+/NLucTom^* NSCs treated with (dotted green and orange lines, respectively) or without (gray and black, respectively) 5-Aza for 3 days.

Previous work has also shown that 5-Aza treatment in combination with *Xist* knockdown results in X chromosome reactivation in MEFs (8). Therefore, we performed a similar analysis on our *Xist^-/+^:Mecp2^+/NLucTom^* mNSCs. Treatment of cells with 0.5 μM 5-Aza for three days resulted in a significant 10-fold upregulation of NanoLuciferase activity (Fig. 3*B*). If *Xist* was knocked down with ASOs in combination with higher amounts of 5-Aza, reactivation was synergistic and 100-fold higher compared to *Xist* knockdown only, or much higher compared to the background of untreated cells (Fig. 3*B*, Fig. S3*A*). Nevertheless, this reactivation still represented around 0.5-1% of *Mecp2*-NanoLuciferase expression from an active X chromosome in homozygous *Mecp2^NLucTom/NLucTom^* NSCs.

While NanoLuciferase bioluminescent analysis is performed at a populational level, flow cytometry allows us to distinguish Tomato fluorescence at the single-cell level. FACS analysis showed that the entire population of cells shifts towards increased Tomato expression after 10 μM 5-Aza treatment for three days (Fig. 3*C*), irrespective of whether *Xist* is knocked down or not. This disagrees with the fact that *Xist* knockdown and 10 μM 5-Aza treated cells show a 100-fold increase in NanoLuciferase activity compared to 10 μM 5-Aza-only treated cells (Fig. 3*B*), suggesting that NanoLuciferase bioluminescence is more sensitive than fluorescence. The shift of the entire population after 5-Aza treatment towards higher Tomato is due to autofluorescence since WT female cells equally treated with 5-Aza also show indistinguishable increased Tomato fluorescence (Fig. 3*D*). We noticed however a small shoulder on the Tomato-High side of the *Xist* ASO plus 5-Aza-treated population. We then proceeded to FACS sort the Tomato-Low, -Medium and -High populations of *Xist^-/+^:Mecp2^+/NLucTom^* mNSCs treated with *Xist* ASOs and 5-Aza and subsequently performed RNA-seq, along with control ASO and non-5-Aza-treated non-FACS-sorted cells (Control). Among the 2,612 genes on the X chromosome, we obtained sufficient allelic expression information from 447 active genes of which 45 were classified as escapees, such as previously described *Mid1, Eif2s3x, Kdm5c, Ddx3x*, etc (Fig. S3*B* and *C*; Yang 2010, Berletch 2015). As expected, *Xist* was expressed from the cast Xi and had decreased expression after its knockdown (Fig. S3*D*). Allele-specific differential expression (DE) analysis showed that 86 genes became reactivated from the cast Xi in the Tomato-High population upon *Xist* knockdown and 5-Aza treatment, *Mecp2* included (Fig. 4*A*, Fig. S3*D*). Reactivated genes were seemingly located in a random fashion along the X chromosome although several clusters were observed (Fig. 4*B*). Among these 86 genes, 7 were more significantly reactivated than *Mecp2* (Fig. 4*A*, Table S2). In addition, *Mecp2* reactivation in the Tomato-Low and -Medium populations was not significant, as expected from the FACS analysis indicating this population to reflect autofluorescence. However, most of the genes within the 86 gene pool in the Tomato-High population were readily reactivated in the low- and medium-Tomato populations (in black, Fig. S3*E*, Table S2), while several genes only significantly reactivated in the Tomato-High population as *Mecp2*. This again suggests that reactivation by *Xist* knockdown and a DNMT1 inhibitor happens more readily for other genes than *Mecp2*.

**Fig. 4.**
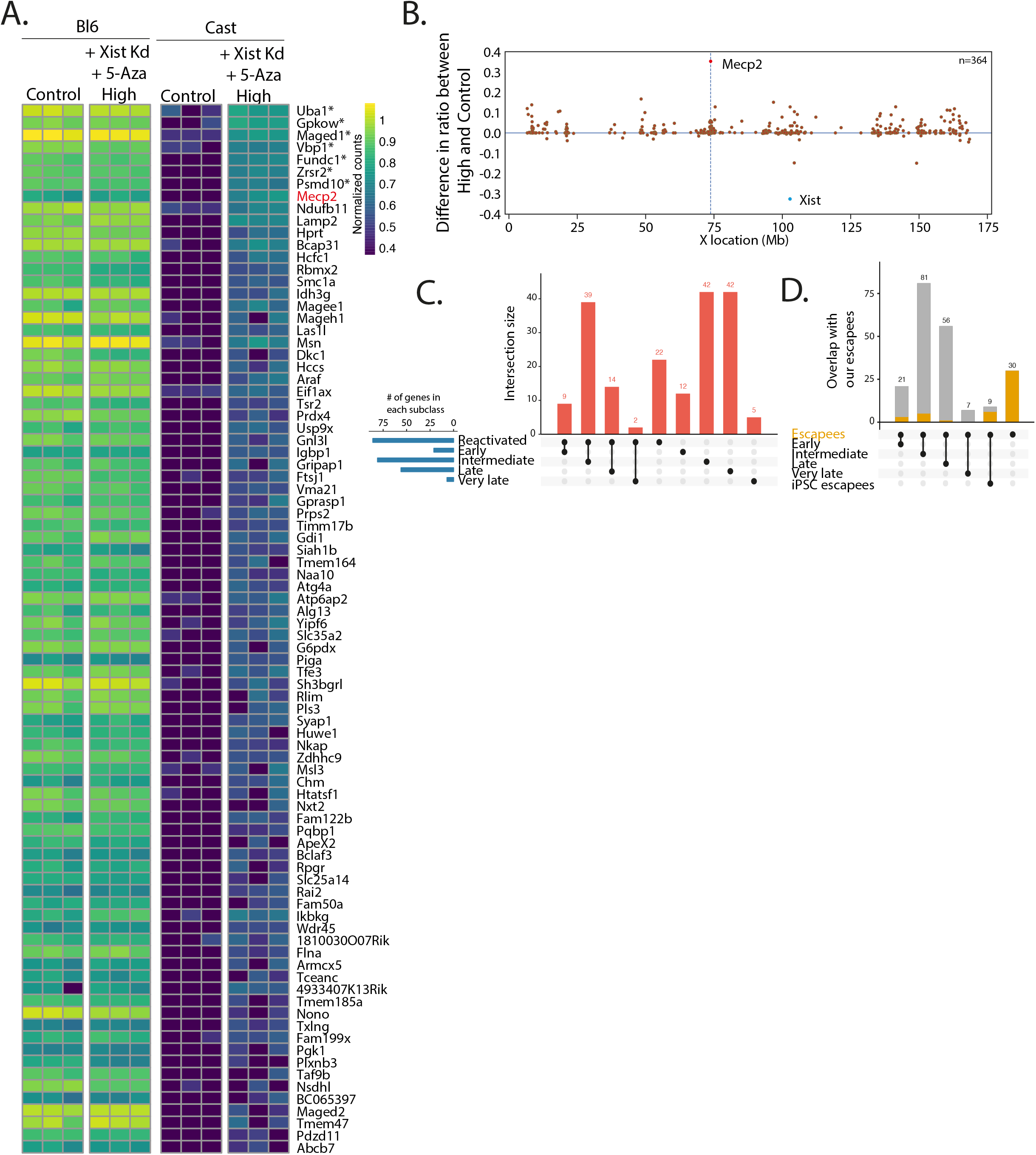
Reactivation of many X-linked genes after *Xist* knockdown and 5-Aza treatment. (***A***) Heatmap showing the normalized allele-specific counts for the 86 genes that are significantly reactivated from the cast Xi in the FACS-sorted Tomato-High population compared to the negative control sample. *Mecp2* is colored in red. The seven genes more easily reactivated than *Mecp2* bear an asterisk. (***B***) Difference between the ratios of cast expression to total expression (Bl6 + cast) of high and control per gene along the X chromosome. Only genes with sufficient allele-specific reads in both conditions are shown. (***C***) Upset plot showing the overlap between our reactivated genes and the early, intermediate, late and very late gene subclasses in (10). (***D***) Upset plot showing the overlap between our escapees and the early, intermediate, late, very late and escape gene subclasses in iPSCs reprogramming (10). For each iPSC gene class, the number of overlapping genes with our escapees and non-overlapping genes are shown in orange and gray, respectively.

In a previous study of iPSC reprogramming of female MEFs, the authors describe different X-linked gene subclasses based on their X chromosome reactivation kinetics, namely early, intermediate, late and very late reactivation (10). We compared our pool of reactivated genes with theirs and observed that 9 of our genes were among the 21 early reactivated iPSC genes (43%), implying that 12 of their early genes were not reactivated in this study (Fig. 4*C*). Similarly, 39/81 (48%), 14/56 (25%) and 2/7 (29%) of our genes were found in their different intermediate, late and very late inactivation gene subclasses, respectively. This means that a low number of our reactivated genes (22/86, 26%) were not reactivated in the iPSC study. Of note, notwithstanding NSCs are very different from reprogramming MEFs, 6 out of their 9 escapees are among our escapee gene pool (67%), while only 9 of their 165 inactivated genes (5%) were in our escapee gene list (Fig. 4*D*), suggesting that our escape genes are not spuriously reactivated genes due to culture conditions for instance.

### Genomic and epigenomic features of X chromosome reactivation

Since our NSCs were subject to 5-Aza treatment, we investigated whether gene reactivation is dependent on CpG-methylation loss. We firstly analysed the density of CpGs in the reactivated and non-reactivated subclasses and found that reactivated genes have significantly more CpGs near their TSS than non-reactivated genes, but did not show differences in CpG density in their gene bodies (Fig. 5*A*, Fig. S4*A*). We subsequently performed MeD-seq analysis (12) on the Control and Tomato-High populations to assess the methylation status on the Xi. We additionally analysed male WT NSCs to assess the methylation status of CpGs on the Xa, and in this way be able to infer CpG methylation on the Xi of our female cells. We observed a global decrease in methylation on the X chromosome as expected from the 5-Aza treatment (Fig. S4*B*). However, we surprisingly could not find any correlation between loss of CpG methylation and reactivation of genes on the Xi. In fact, the promoter of *Mecp2* and other X-linked genes were methylated in control NSCs. While male NSCs showed a complete lack of DNA methylation as expected from expressed genes, promoters in female cells were not significantly demethylated in the Tomato-High population compared to control NSCs, although methylation seemed lower (Fig. S4*C*, Table S3). However, 16 out of our 86 reactivated genes showed significant lower levels of DNA methylation (Fig. S4*C*, Table S3). We performed a cluster analysis of the CpG methylation status of promoters of both reactivated as non-reactivated genes and observed no clustering of reactivated promoters (Fig. S4*D*). Altogether, loss of methylation was not a clear indicator of X-linked gene reactivation, pointing to other mechanisms at play.

**Fig. 5.**
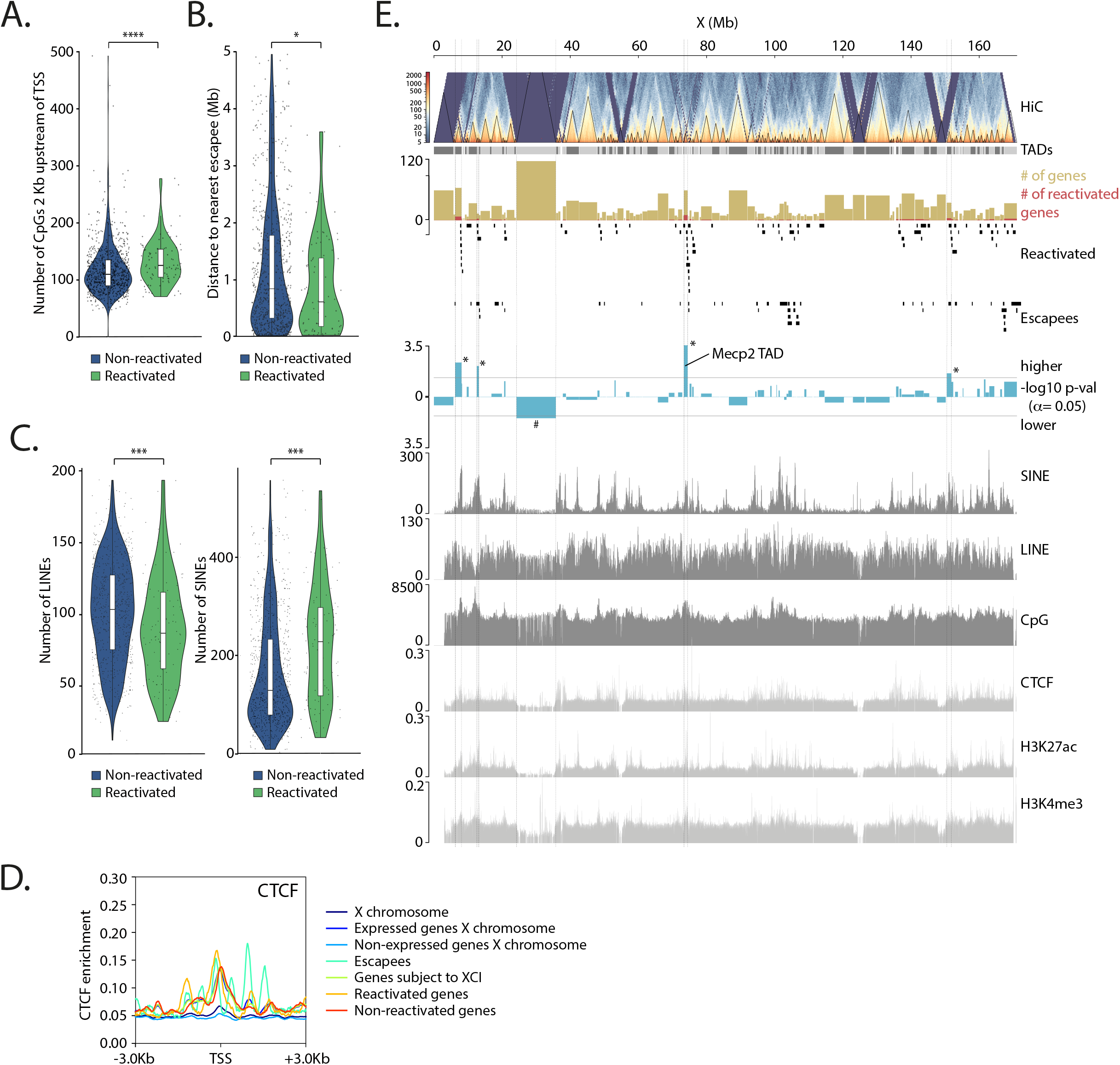
Reactivation of many X-linked genes correlates with genomic and epigenomic features. (***A***) Violin plot depicting the number of CpGs in a bin of 2 Kb upstream of the TSS of non-reactivated (blue) and reactivated genes (green). (***B***) Violin plots of the distance to the nearest escapee in Mb of non-reactivated (blue) and reactivated genes (green). (***C***) Violin plots of the number of LINEs and SINEs in a window of ± 200 Kb around the TSSs of non-reactivated genes (blue) and reactivated genes (green). * p-val<0.05, ** p-val<0.01, *** p-val<0.001, **** p-val<0.0001, Mann-Whitney test. (***D***) Density plots of CTCF binding to the TSS ± 3 Kb of different gene subclasses. (***E***) Genome browser overview showing TADs overlapping the X-chromosome. NPC Hi-C data from Bonev et al. 2017 is shown from which the TADs were identified. For each TAD, the number of overlapping reactivated genes and escapees were identified, as plotted here along the X-chromosome. The yellow and red rectangles show the number of genes and reactivated genes per TAD, respectively. The –log10 of the p-value of a Binomial test between the ratio of reactivated genes / total genes per TAD and for the whole X-chromosome are shown in blue for each TAD. TADs with a higher or lower ratio than on average are plotted in inverse directions. A p-value of 0.05 is indicated by a dotted line, and TADs with a significantly higher and lower ratio are depicted with an * and #, respectively. SINE, LINE, CpG, CTCF, H3K27ac and H3K4me3 densities along the X chromosome are depicted in gray underneath.

We subsequently performed genomic feature correlation analyses on our list of reactivated genes. Firstly, we did not detect a correlation between the position of reactivated genes on the X and proximity to *Xist*, as previously described for X-linked reactivated genes during iPSC reprogramming (10)(Fig. S4*E*). However, genes that are reactivated tend to be closer to escapees than non-reactivated genes (Fig. 5*B*), suggesting that proximity to an escapee is a determining factor in the reactivation potential of X-linked genes. In addition, we find that genes that are more easily reactivated tend to have significantly less LINEs and more SINEs around their TSSs (Fig. 5*C*). Genes that are significantly reactivated tend to have higher numbers of any SINE subclasses compared to non-reactivated genes (Fig. S4*F*).

We next investigated the correlation between our different gene subclasses with a published CTCF binding profile and several chromatin marks ChIP-seq datasets obtained from ESC-derived male neural progenitor cells bearing thus a single Xa (Bonev 2017). Escapees tend to show increased enrichment of CTCF at their TSSs (Fig. 5*D*), as has been previously described (13, 14). In addition, our reactivated genes tend to have more CTCF binding at their TSSs on the Xa compared to non-reactivated genes, while also bearing increased H3K4me3 and H3K27ac deposition (Fig. 5*D*, Fig. S4*G*).

We furthermore examined whether certain TADs are more easily reactivated or prevented from reactivating than others by crossing our gene subclasses with previously published TAD data from male NPCs (Bonev 2017). Based on the number of reactivated genes within each TAD, we identified 4 TADs with significantly more reactivated genes compared to the whole X chromosome (as indicated by an *) (Fig. 5*E*). The TAD containing *Mecp2* shows the significantly largest ratio of reactivated genes, probably due to the fact that the RNA-seq analysis was performed on Tomato-High (*Mecp2*-reactivated) sorted cells. Nevertheless, these four TADs contain a low number of escapees compared to other non-significant TADs (Fig. 5*E*). Interestingly, a single TAD is particularly resistant to reactivation (as indicated by a #) and yet, contains many genes. We subsequently investigated whether the presence of CpGs, SINEs and LINEs within those 5 TADs could be indicators of their tendency or resistance to reactivate. TADs with significantly more reactivated genes than other TADs tended to have more CpGs and less LINEs, (although not significant) and contained significantly more SINEs (Fig. S4*H*).

## Discussion

A proper mouse model to study reactivation of *Mecp2* from the Xi in a very sensitive manner has been lacking. We provide here a new mouse model where *Mecp2* has been fused with the bioluminescent reporter NanoLuciferase, which is a 100-times brighter than the frequently used firefly luciferase, and a fluorescent reporter Tomato. This dual capability not only permits measurement of NanoLuciferase activity at a populational level, but also measurement of Tomato fluorescence at the single-cell level and *in vivo. Mecp2^NLucTom^* mice are viable and have been created in a Cast/EiJ background which allows tracking the level of reactivation in a chromosome-X- and genome-wide manner thanks to the presence of hundreds of thousands of informative SNPs with respect to the more commonly used C57BL/6 and 129/Sv strains.

By using B6 females carrying an oocyte-specific *Zp3-Cre* transgene and a *Xist^2lox^* allele, we have generated a maternal knockout of *Xist*. Crossing these females with *Mecp2^NLucTom^* cast males has allowed us to generate *Xist^-/+^:Mecp2^+/NLucTom^* embryos. An alternative model where the females carry the *Zp3-Cre, Xist^2lox^* and *Mecp2^2lox^* alleles has allowed us to generate RTT-affected *Xist^-/+^:Mecp2~^-/NLucTom^* embryos. We have derived ESCs, MEFs and NSCs from these F1 embryos. *Xist^-/+^:Mecp2^+/NLucTom^* and *Xist^-/+^:Mecp2^-/NLucTom^* NSCs show skewed XCI as expected by the presence of the *Xist* deletion on the maternal B6 X chromosome while not showing any *in vitro* escape of *Mecp2* from the Xi. Why our *Xist^-/+^:Mecp2:^-/NLucTom^* neurons do not show RTT-related phenotypes is unclear. Most RTT-affected neuronal studies have been performed with ex vivo neuronal cultures (15–17). However, *Mecp2* knockout neurons obtained by ESC differentiation showed smaller nuclear size than WT neurons after long-term culture in vitro (18). It is thus possible that our 10-11-day NSC differentiation is not sufficient to bring RTT phenotypes to the fore.

We have tested several compounds to assess whether the reporters can be reactivated. Contrary to what has been previously published (7), neither individual or combined treatments with LDN193189, GSK650394 or RG2833 resulted in *Mecp2* reactivation in our *Xist ^-/+^:Mecp2 ^-/NLucTom^* NSCs. These differences might be due to Przanowski and colleagues using fibroblasts and adult brains instead of NSCs, or our NSCs might be more resilient to reactivation. In addition, another inhibitor of ACVR1, K0228, also failed to reactivate a *Mecp2*-Luciferase reporter in mouse tail fibroblasts (19). However, combined treatment of GSK650394, LDN193189 and 5-Azacitidine resulted in similar reactivation of *Mecp2* compared to 5-Aza only.

We have synergistically reactivated our NanoLuciferase-Tomato reporter with a combined treatment of 5-Aza and *Xist* knockdown. FACS analysis showed that a small population of treated cells shifted towards high Tomato fluorescence, while RNA-seq analysis indicated that a substantial population of cells in this fraction respond to the treatment and reactivate *Mecp2*, although, as expected, reactivation is not *Mecp2*-specific. 85 additional genes become significantly reactivated and several amongst these are more easily reactivated than *Mecp2*, which is something that will have to be taken into consideration when using general XCR methods with drugs as therapeutic treatments. Strikingly, we observed a significant overlap between our reactivated gene pool and genes reactivated early and intermediate late by means of IPSC reprogramming of female MEFs (10). We therefore conclude that many X-linked genes show a predisposition to reactivate regardless of the technique, be it *Xist* knockdown combined with 5-Aza treatment, or overexpression of the OCT4, SOX2, KLF4, MYC transcription factors.

We examined which genetic or epigenetic mechanisms leading to XCR are at play here. Correlation analysis of reactivated genes with CpG presence and methylation loss after 5-Aza treatment indicate that although increased CpG presence is an indicator of reactivated genes, their reactivation surprisingly does not seem to be dependent on methylation loss. However, this can be reconciled with the fact that a small reduction in promoter methylation, not detectable by MeD-seq, might be sufficient for gene re-expression. This may also explain why we only detect limited reactivation of *Mecp2* by NanoLuciferase activity analysis. In addition, decreased distance to escapees while increased SINE and decreased LINE densities are potent indicators of reactivation. Correlating with our study, genes that are more easily silenced on the X chromosome or are ectopically silenced on chromosome 12 tend to have more LINEs and less SINEs close to their TSSs (13). Moreover, in line with our results, X-linked genes that are early reactivated during iPSC reprogramming of female MEFs harbor increased number of SINEs closer to them than late or very late reactivating genes (10). There are strong indications that SINEs and LINES may play an important role in the capability of genes to be silenced or reactivated. SINE-mediated expansion of CTCF binding sites might explain why we detect an Xa-specific increased binding of CTCF and an increased number of SINEs closer to reactivated genes (20, 21). Nevertheless, reactivated genes show an enrichment of all subclasses of SINEs irrespective of their age, and not only CTCF-enriched SINE B2 transposable elements (21). Hence, how SINE elements might be involved in silencing and reactivation of X-linked genes remains an open question. SINEs may be involved in setting up higher order chromatin structure to overcome gene repression. In addition, two histone marks of promoter activity on the Xa, H3K4me3 and H3K27ac, also tend to correlate with XCR, indicating that genes with strong activity signatures on the Xa are more easily reactivated, probably due to their higher capacity to attract transcription factors.

Finally, we interrogated proclivity of X-linked TAD to reactivate. Likely because we selected a reactivated population based on Tomato fluorescence, we find that the TAD containing *Mecp2* is more easily reactivated than other TADs. Three other TADs also show a tendency to more easily reactivate than other TADs. Their tendency to reactivate correlates again with a higher presence of SINEs, in line with our results showing SINEs to be strong indicators of reactivation potential. In conclusion, genes that are reactivated by *Xist* knockdown and 5-Aza treatment overlap significantly with genes that are reactivated by other means, namely during reprogramming of MEFs towards iPSCs, suggesting general intersecting mechanisms for XCR.

Altogether, we describe here a new mouse model system which is more sensitive than any bioluminescent and fluorescent system currently available in the community to study reactivation of Mecp2 to treat Rett Syndrome, *in vitro* and *in vivo*. These mouse lines could be used to study Mecp2 reactivation by high-throughput screening of chemical compounds or by more targeted approaches such as CRISPR/Cas9 fused to activators or repressors.

## Acknowledgements

We thank Esther Sleddens-Linkels and Tsung Wai Kan, from the Developmental Biology and Biochemistry Departments, Erasmus MC, for help with cell sorting. We thank Martí Quevedo and Raymond Poot from the Cellular Biology Department, Erasmus MC, for help with mouse NSC isolation and growth, and comments on the manuscript. We also thank Robin Adrianse and Smitha Sripathi from the Clinical Research Division at the Fred Hutchinson Cancer Research Center for kindly offering plasmid information and 5-Aza advice. We thank Laura Mezzanotte and Yanto Ridwan form the Radiology and Nuclear Medicine Department, Erasmus MC, for helping with furimazine injections and imaging. Finally, we thank Maurice de Wit from the Department of Neurology, Erasmus MC, for helping with taking pictures of neural cultures and neurite analysis. This research was funded by the Rett Syndrome Research Trust.

## Figures

**Fig. S1.**
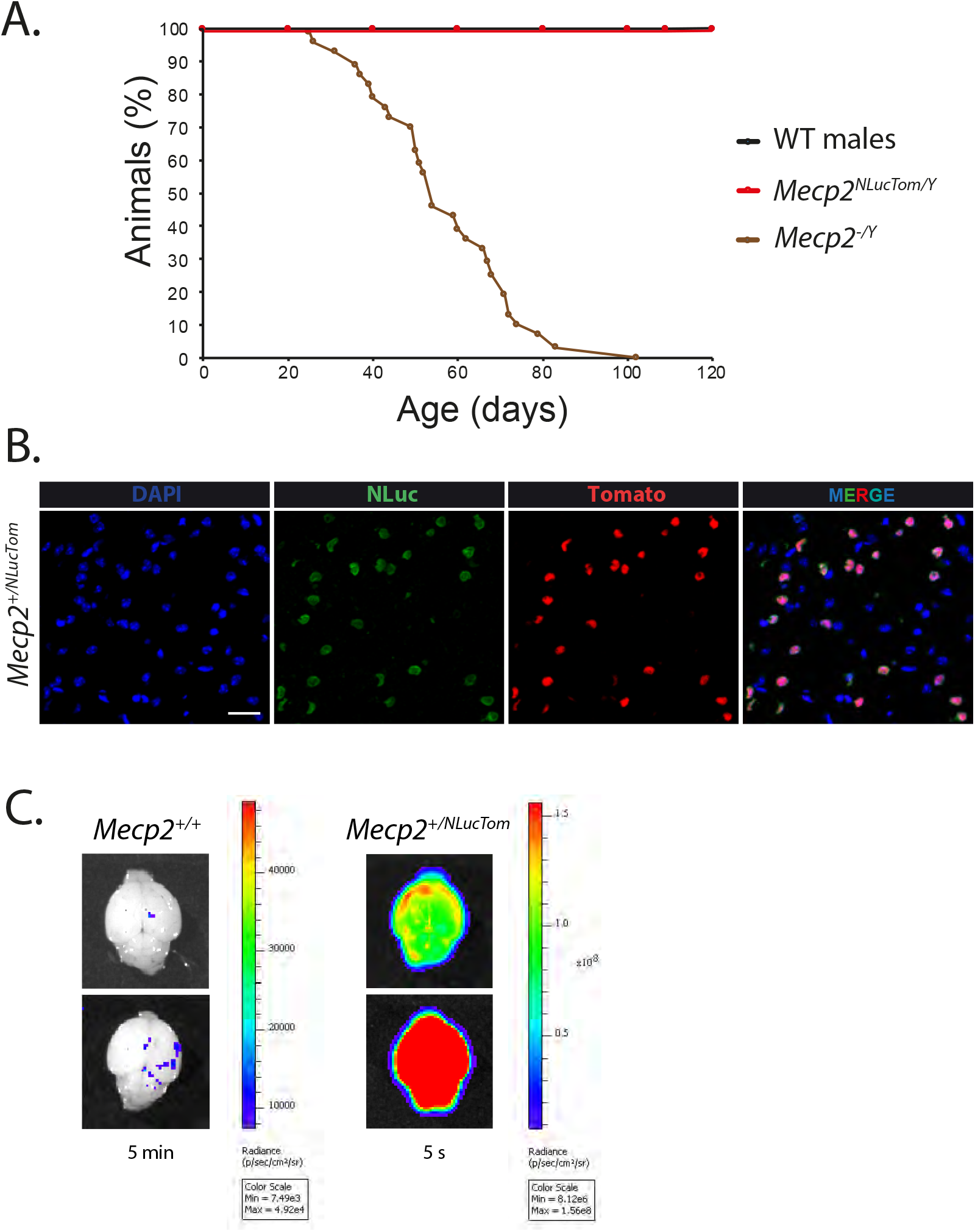
Cast *Mecp2* NLucTom mice are viable and express the fusion protein in the brain. (***A***) Cumulative survival plot showing the % of cast *Mecp2^NLucTom/Y^* males (red) compared to WT cast males and *Mecp2^Δ/Y^* males (22) surviving at a given time in days. (***B***) IF of NanoLuciferase (green) and endogenous Tomato fluorescence (red) in brain sections of heterozygote *Mecp2^+/NLucTom^* females showing random XCI. DAPI, blue. White scale bar: 25 μm. (***C***) Representative bioluminescence images of two P6 WT female brains and two P6 *Mecp2^+/NLucTom^* female brains after furimazine injection.

**Fig. S2.**
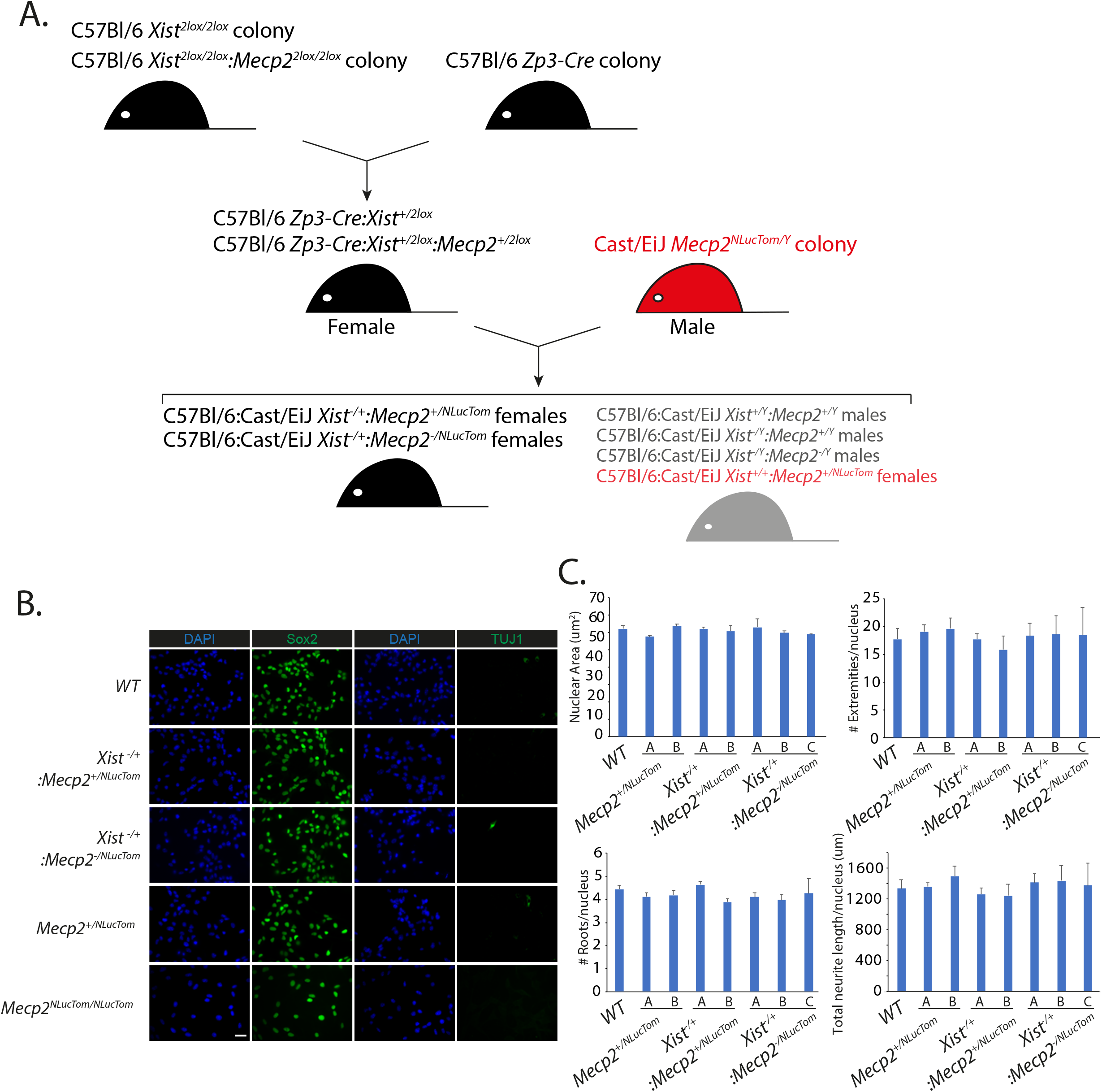
Generation of WT, *Xist^-/+^:Mecp2^+/NLucTom^, Xist^-/+^:MecpZ^-/NLucTom^, Mecp2^+/NLucTom^* and *Mecp2^NLucTom/NLucTom^* NSCs. (***A***) Breeding strategy to generate *Xist^-/+^:Mecp2^+/NLucTom^*, *Xist^-/+^:Mecp2^-/NLucTom^* embryos and associated lines. We keep four independent colonies in two backgrounds: C57BL/6 *Xist^2lox/2lox^*, C57BL/6 *Xist^2lox/2lox^:Mecp2^2lox/2lox^*, C57BL/6 *Zp3-Cre* and Cast/EiJ *Mecp2^NLucTom^* mice. *Xist^2lox/2lox^* and *Xist^2lox/2lox^:Mecp2^2lox/2lox^* females or males are crossed with *Zp3-Cre* mice to generate heterozygous *Zp3-Cre:Xist^+/2lox^* and *Zp3Cre:Xist^+/2lox^:Mecp2^+/2lox^* females in a C57BL/6 background. These females are then crossed with *Mecp2^NLucTom/Y^* males, to generate hybrid *Xist ^-/+^:Mecp2^+/NLucTom^* or *Xist^-/+^:Mecp2:^/NLucTom^* female embryos. Other possible genotypes of this last crossing are depicted in gray. Notice all genotypes from this second crossing can have Zp3-Cre (50% probability), not depicted in figure. (***B***) IF of Sox2 or TUJ1 (left and right green respectively) in WT, *Xist^-/+^:Mecp2^+/NLucTom^, Xist^-/+^:Mecp2^YNLucTom^, Mecp2^+/NLucTom^* and *Mecp2^NLucTom/NLucTom^* NSCs. DAPI, blue. White scale bar: 25 μm. (***C***) Nuclear area, number of cellular extremities per nucleus and number of roots per nucleus (average ± s.d., n=3 biological replicates, 191-521 neurons per replicate) of 2-3 independent WT, *Xist^-/+^:Mecp2^+/NLucTom^* and *Xist^-/+^:Mecp2^-/NLucTom^* NSC clones.

**Fig. S3.**
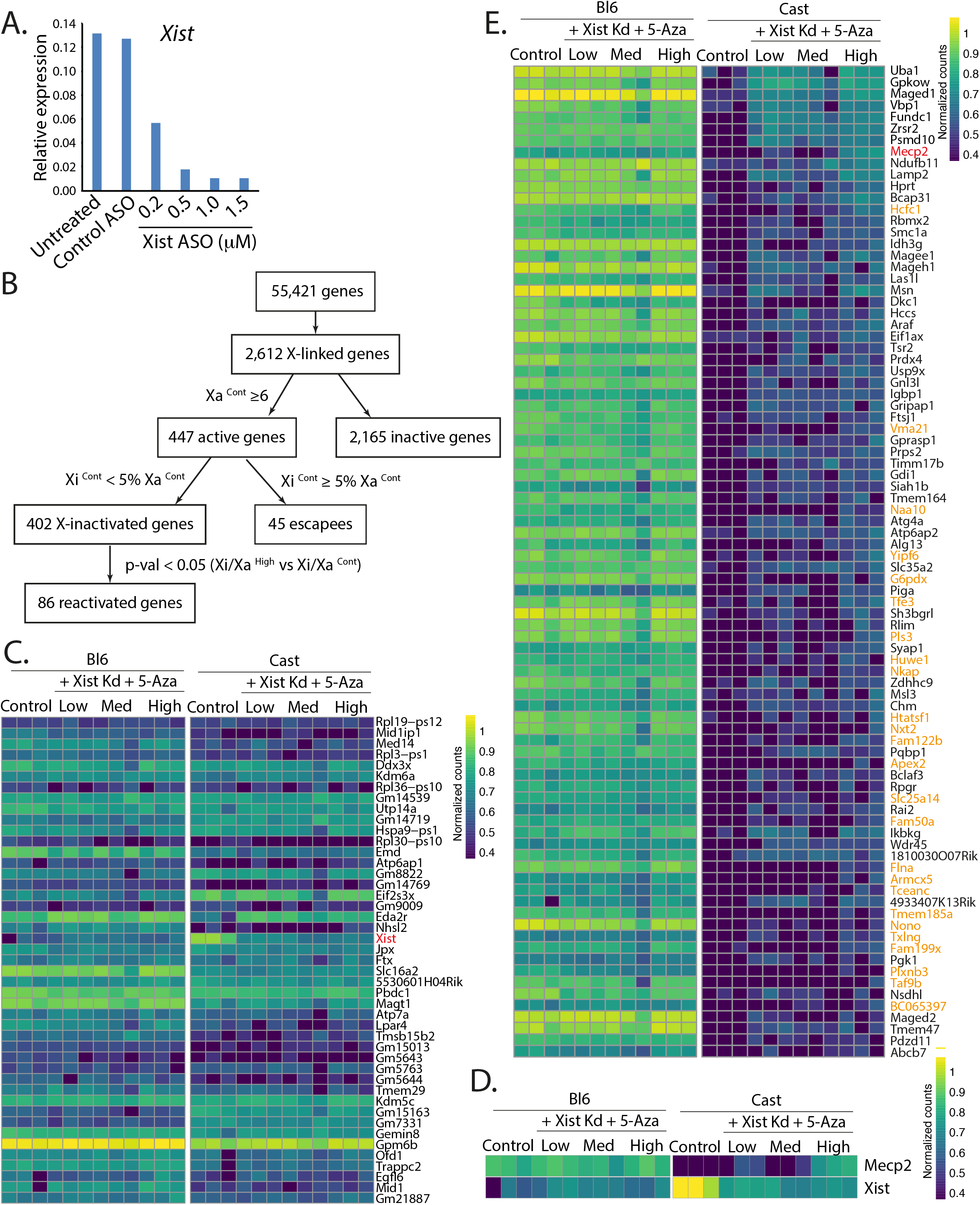
Many X-linked genes are reactivated in mNSCs after *Xist* knockdown and 5-Aza treatment. (***A***) Relative *Xist* expression in *Xist^-/+^:Mecp2^+/NLucTom^* NSCs after knockdown of *Xist*with *Xist* ASOs or control ASOs. Different concentrations of *Xist* ASOs were tested, n=1. (***B***) Flowchart indicating the general steps performed to obtain the different gene subclasses from the RNA-seq analysis. (***C***) Expression heatmap of the different escapees across the different samples and alleles. *Xist* is indicated in red. (***D***) Expression heatmap of *Mecp2* and *Xist* across the different samples and alleles showing *Mecp2* reactivation in the Tomato-High population and *Xist*downregulation in the Tomato-Low, -Med and -High populations as expected from the knockdown experiment. (***E***) Expression heatmap of the reactivated genes in the Tomato-High population across the different samples and alleles, ordered by P-value. *Mecp2* is indicated in red. Genes that are also reactivated in the Tomato-Low and/or Tomato-Medium populations are indicated in black, while genes only significantly reactivated in the Tomato-High population as *Mecp2* are indicated in orange.

**Fig. S4.**
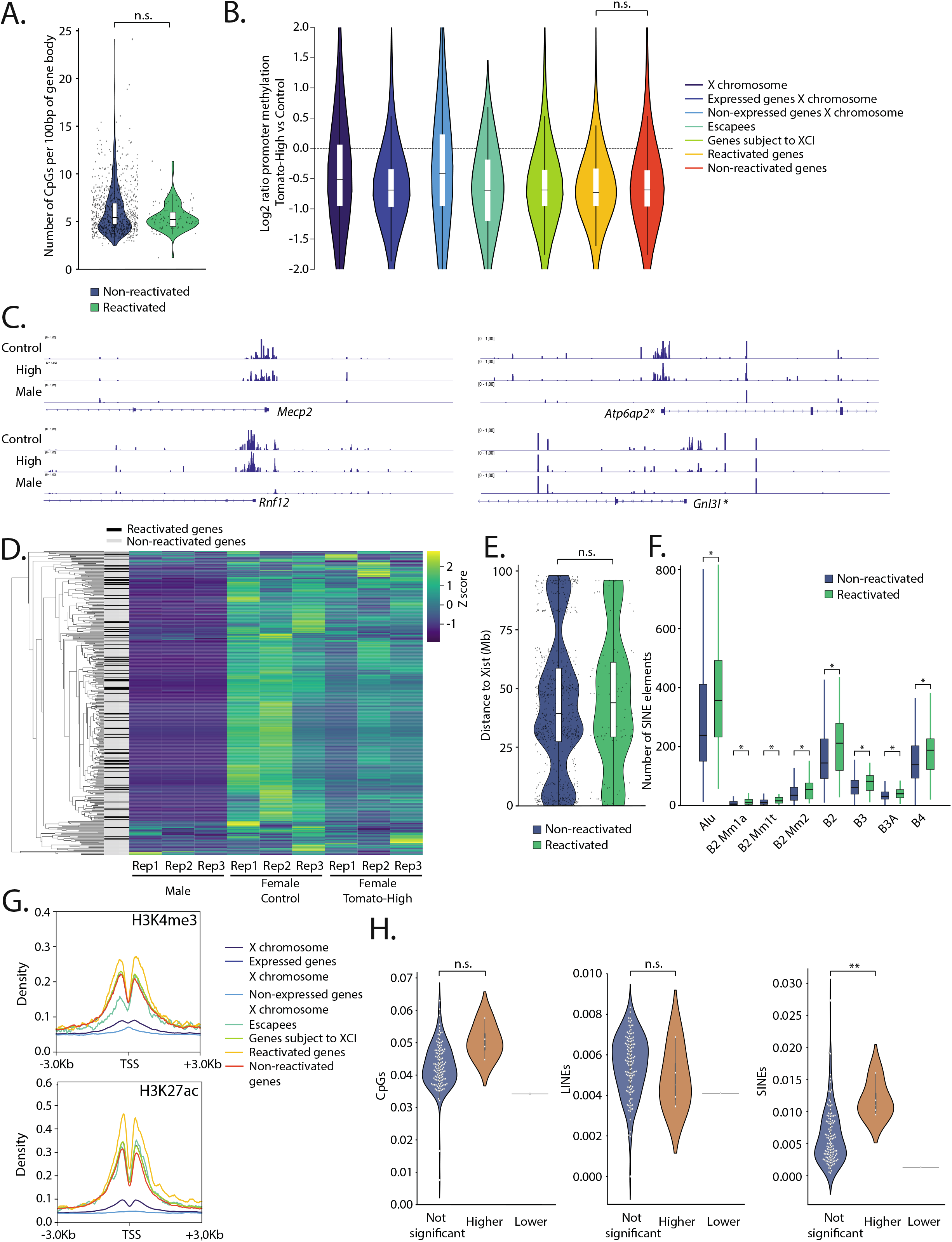
Reactivation does not correlate with a general loss of methylation at promoters. (***A***) Violin plot depicting the number of CpGs per 100bp of gene body of reactivated and non-reactivated genes; n.s., not significant (Mann-Whitney test, p-value < 0.05). (***B***) Violin plots depicting the log2 ratio between the methylation status of the promoter (± 1 Kb around TSS) of the different gene subclasses in the Tomato-High and control populations. Dotted line indicates an identical methylation status for both conditions (ratio = 1); n.s., not significant (Mann-Whitney test, p-value < 0.05). (***C***) Genome browser overview with the average normalized MeD-seq tracks at the promoter areas of four reactivated genes *Mecp2, Rnf12, Atp6ap2* and *Gnl3l* in female control, female Tomato-High and male NSCs. Genes with significant loss of DNA methylation at their promoters (±1 Kb of the TSS) are indicated by an asterisk. (***D***)) Heatmap of DNA methylation status around the TSSs of reactivated genes and non-reactivated genes of the three biological replicates of female control, female Tomato-High and male NSCs. Z-scores of MeD-seq read counts ±1 Kb of the TSS are shown. Next to the clustering dendrogram, genes are annotated as reactivated and not-reactivated in black and gray, respectively. (***E***) Violin plot depicting the distance to *Xist* in Mb of non-reactivated genes (blue) and reactivated genes (green). Note the non-significant difference between both populations; n.s., not significant (Mann-Whitney test, p-value < 0.05). (***F***) Boxplots depicting the number of different SINE subclasses in ± 200 Kb bins around the TSS of non-reactivated genes (blue) and reactivated genes (green). * p-val<0.05, Mann-Whitney test. (***G***) Average density plots showing the H3K4me3 and H3K27ac enrichment in the ± 3 Kb region around the TSS of the different gene subclasses. (***H***) Violin plots of the normalized number of CpGs, LINEs and SINEs in TADs that are not significantly enriched for reactivated genes and TADs that are significantly enriched (higher) or depleted (lower) for reactivated genes. * p-val<0.01, Mann-Whitney test, n.s. not significant.

---

Table S1. Primers used in this study.

Table S2. X-linked differential expression analysis between the Tomato-High and Control populations.

This table shows the differentially expressed X-linked genes between the Tomato-High and Control populations using DESeq2. Genes are annotated according to their gene group and X-inactivated genes that are reactivated in the Low and Medium samples are indicated.

Table S3. DNA methylation analysis of X-linked promoters using MeD-seq.

This table shows the normalized MeD-seq reads of male, Tomato-High and Control populations at the X-linked gene promoters (± 1kb of the TSS). Fold change between Tomato-High and Control are reported, as well as the p-value based on a Mann-Whitney test. Genes are annotated according to their gene class based on the RNA-seq data.

## Materials and Methods

### Mouse lines and work

All animal experiments were performed according to the legislation of the Erasmus MC Rotterdam Animal Experimental Commission.

*Mecp2^NLucTom^* mice were obtained from *Mecp2^NLucTom^* ESCs generated in house (see below) and kept in a cast background colony. *Xist^2lox^* mice (23) were kept in a B6 background. *Mecp2^2lox^* and *Zp3-Cre* mice were obtained from the Jackson Laboratory (B6.129P2-MeCP2tm1Bird/J - #077177, C57BL/6-Tg(Zp3-Cre)93Kw/J - #003651 respectively) and kept in a B6 background. *Xist^2lox^* mice were crossed with *Mecp2^2lox^* mice to generate a colony of *Xist^2lox/2lox^:Mecp2^2lox/2lox^* mice. *Xist^2lox/2lox^* and *Xist^2lox/2lox^:Mecp2^2lox/2lox^* female mice were crossed with male *Zp3-Cre* mice to generate *Xist^2lox/+^ Zp3-Cre* and *Xist^2lox/+^:Mecp2^2lox/+^ Zp3-Cre* females. These were then crossed with cast *Mecp2^NLucTom/Y^* males to generate *Xist^-/+^:Mecp2^+/NLucTom^, Xist ^-/+^:Mecp2-^-/NLucTom^* and *Xist^+/+^:Mecp2^+/NLucTom^* female hybrid embryos. *Mecp2^NLucTom/NLucTom^* female embryos were obtained from the *Mecp2^NLucTom^* cast colony. WT hybrid females were obtained by crossing B6 females with cast males.

### Cell culture

All ESC lines were grown in regular ESC medium (DMEM, 10% fetal calf serum, 100 U ml^-1^ penicillin/streptomycin, 0.1 mM 2-mercapoethanol, 0.1 mM NEAA, 5000 U mL^-1^ LIF) supplemented with 2i (1 μM PD0325901 – Selleckchem, 3 μM CHIR99021 – Axon Medchem) on irradiated male mouse embryonic fibroblasts.

ESCs were generated from E3.5 blastocysts. Briefly, E3.5 blastocysts were flushed from uteri in M2 medium. Their zona pellucida was removed with acidic Tyrode’s solution (Sigma) at RT for several seconds. Embryos were subsequently washed in M2 medium and transferred to 4-well plates, one blastocyst per well, containing irradiated MEFs and regular ESC medium supplemented twice as much as the normal amount of LIF and PD98059 (final concentration 50 μM; Cell Signaling Technologies). ICM outgrowths were then picked 5-7 days plating the blastocyst and expanded. Once ESCs reached the 12 well stage, they were genotyped. The selected genotypes were grown as single cells/colonies by plating them in serial dilutions in 10 cm dishes. One day after plating, cells were weaned off the increased concentration of LIF and of PD98059. Colonies were picked, selected for morphology and correct karyotype.

To target *Mecp2* in male cast ESC cells, 0.6×10^°6^ ESCs were transfected with 2 μg CRISPR/Cas9 targeting the *Mecp2* STOP codon and 2 μg of the donor vector carrying 5’ and 3’ 500bp-long cast-specific homology arms with a NanoLuciferase and Tomato reporters in frame with *Mecp2’s* coding region. A P2A signal between NanoLuciferase and Tomato leads to Tomato being translated as an independent protein. ESCs were transfected with 4 μL lipofectamine 2000 following the manufacturer’s instructions (ThermoFisher). 2 days after transfection, Tomato-positive cells were FACS-sorted, put back into culture for a few days and subsequently injected in B6 blastocysts.

MEFs were isolated from E12.5 embryos. E12.5 embryos were removed from yolk sacs and their heads, liver, hearts and digestive tract were removed. The remaining carcass was chopped into fine pieces and added to 5 mL of Trypsin-EDTA (Life Technologies), for 10 min 37°C in water batch. Falcons were shaken smoothly every 2 min. Remaining clusters and cells were pipetted up and down several times. Medium was added to quench the trypsin and the sample was centrifuged 5 min 1000 rpm. Cell pellet was resuspended in regular ESC medium without LIF and 2i and grown in 0.2% gelatin-coated 15cm dishes. After 3x 15 cm dishes were obtained, cell lines were genotyped and frozen.

NSC lines were isolated from E15.5 embryos. E15.5 brains were extracted, hemispheres were cut and meninges dissected when possible. Cortexes were chopped into pieces and introduced in 15 mL falcon tubes containing 2.5 mL dissecting medium (PBS +3% glucose) plus 300 μL Trypsin-EDTA (10x; Life Technologies, 15400-054). The falcon was then incubated for 10 min at 37°C while shaking it every 2 min manually. Trypsin was then inactivated with 500 μL horse serum (Life Technologies, 16050-130). 50 μL DNaseI (1mg mL^-1^) was added and incubated for 8-10 min at 37°C. Pipette up and down around 10 times with 1 mL pipette. Centrifuge 1000 rpm 5 min and resuspend in filtered NSC culture medium (Conti 2005): 192 mL EuroMed-N (EuroClone), 2 mL N2 supplement (Invitrogen), 1 mL human insulin (final concentration 20 μg mL^-1^, Roche), 1ml BSA (final concentration 50 μg mL^-1^, Gibco), 2 mL L-Glut (100x; Gibco), 10 μL murine EGF (final concentration 10 ng mL^-1^, Peprotech), 10 μL human bFGF (final concentration 10 ng mL^-1^, Peprotech), 100 U mL^-1^ penicillin/streptomycin. Cells were grown in suspension in 10cm dishes for a week. Resulting neurospheres were disaggregated and cells were grown as a monolayer. The different lines were established and grown in 6-well plates. Wells were precoated with 0.2% gelatin for 5 min RT, then removed and 1.5 mL Laminin (final concentration 5 μg mL^-1^ in PBS; Sigma) was added to wells for at least 5 h at 37°C, better o/n at 37°C, or kept in the fridge with parafilm for up to a month. Prior to use, wells were washed quickly 3x with PBS and cells added. Cells were passaged with Accutase (Sigma) every 3-5 days up to passage 10-12, generating frozen vials along the way.

NSC differentiation into neurons, astrocytes and oligodendrocytes was performed following the protocol by Spiliotopoulos and colleagues (24). NSCs were counted and seeded in 5 mL D1 medium (EuroMed-N, 0.5% N2 (Invitrogen), 1% B27 (Invitrogen), 10 ng mL^-1^ human bFGF and 100 U mL^-1^ penicillin/streptomycin) at 1.35×10^5^ cells cm^-2^ per well of a 6-well dish previously coated with 0.1% gelatin and 5 μg mL^-1^ laminin in PBS for at least 5 hours (day 0). 3 days after plating, cells were collected with accutase, counted and reseeded in 3 mL A medium (1:3 mix DMEM/F12 (Gibco) and Neurobasal (Gibco), 0.5% N2, 1% B27, 10 ng mL^-1^ bFGF, 20 ng mL^-1^ BDNF (Prospec) and 100 U mL^-1^ penicillin/streptomycin) at 5 ×10^4^ cells cm^-2^ per well of a 6-well dish with coverslips precoated with gelatin and laminin (day 3). Medium was changed at day 5 and at day 6, medium was changed by B medium (1:3 mix DMEM/F12 and Neurobasal, 0.5% N2, 1% B27, 6.7 ng mL^-1^ bFGF, 30 ng mL^-1^ BDNF and 100 U mL^-1^ penicillin/streptomycin). At day 9, medium was changed to B1 medium (same as B medium although at 5 ng mL^-1^ bFGF). Coverslips were then fixed at day 10-11 after the start of differentiation following the IF protocol below.

### Immunofluorescence

Mice were perfused with 4% PFA for 3 min. Brains were removed and fixed again in 4% PFA for 1 h at RT. Brains were left to sink o/n in 10% sucrose in PBS at 4°C and frozen in OCT the next day by bathing a freezing cup isopentane in dry ice and stored afterwards at −80°C. 7 μm coupes were generated with a cryostat microtome and put on adhesive slides. Slides were left to dry 30 min at RT and frozen at −80°C. Slides were thawed 30 min at RT and processed with the same IF protocol as cell cultures, see below.

IF on cell cultures were performed as follows. Cells were grown on coverslips and subsequently blocked in 5% goat serum (Sigma, G9023; or donkey serum when necessary, Sigma, D9963), 1% Triton X100 (performed) in PBS for 1 h at RT. The primary antibody was applied in PBS 5% goat (or donkey) serum, 0.1% Triton X1000 0/n at 4°C in the dark. The next day, slides were washed 3x with 5% goat (or donkey) serum, 0.1% Triton X100 in PBS for 5 min at RT. The second antibody was then applied in 5% goat (or donkey) serum, 0.1% Triton X100 in PBS for 1 h at RT and washed 3x with 5% goat (or donkey) serum, 0.1% Triton X100 in PBS for 5 min at RT (last wash containing 1:5000 DAPI). Slides were then mounted with Prolong™ Gold Antifade Mountant (Thermo Scientific). Images were acquired with a fluorescent Axio Imager M2 microscope (Zeiss) and analysed with Fiji and Photoshop software (Adobe). The following primary antibodies were used: mouse anti-TUJ1 (Biolegend, 801202, 1:250), rabbit anti-GFAP (Dako, Z0334, 1:250), rabbit anti-OLIG2 (IBL, 18953, 1:250), goat anti-SOX2 (Santa Cruz, sc-17320, 1:500), rabbit anti-NLuc (kind gift of Promega, 1:100). The following Alexa Fluor secondary antibodies were used: goat anti-rabbit 488 (Invitrogen, A-11008, 1:400), goat anti-mouse 488 (Invitrogen, A-11001, 1:400) and goat anti-rabbit 633 (Invitrogen, A-21070, 1:400), donkey anti-goat 488 (Invitrogen, A-11055, 1:400). IFs of neuronal cultures were not performed with serum, BSA was used instead.

### Western Blot

Cells were harvested in ice-cold PBS with complete protease inhibitors (Roche). Cell pellets were incubated with 400 μL Buffer A (100 mM HEPES, 1.5 mM MgCl2, 10 mM KCl, 0.5 mM DTT and protease inhibitors) for 10 min on ice, vortexed 30 sec and centrifuges 2000 rpm, 5 min, 4°C. Nuclei were then lysed by adding 2x the pellet volumes of Buffer C (20 mM HEPES, 25% glycerol, 420 mM NaCl, 1.5 mM MgCl2, 0.2 mM EDTA, 0.5 mM DTT and protease inhibitors) for 20 min on ice, centrifuged max speed, 2min, 4°C. Protein concentrations were determined with NanoDrop. WB was performed with homemade SDS-PAGE gels and nitrocellulose membranes (Merck). The following antibodies were used, mouse anti-MECP2 (Sigma-Aldrich, M7443, 1:500), rabbit anti-RFP (Abcam, ab62431, 1:500), rabbit anti-NLuc (kind gift of Promega, 1:1000) and b-actin-peroxidase (Sigma, A3854, 1:20,000). Detection of peroxidase activity was performed using ECL western blotting detection reagent (GE Healthcare) in an Amersham Imager 600 (GE Healthcare). Detection of the remaining proteins was performed with an Odyssey CLx imaging system with Imago Studio 5.2 software (LI-COR Biosciences) with the following antibodies: IRDye 800CW donkey anti-mouse and IRDye 680RD donkey anti-rabbit (both from LI-COR Biosciences, 926-32212 and 926-68073 respectively,1:10,000).

### *Xist* knockdown and drug analysis

*Xist* knockdown was performed following manufacturer’s instructions for the Mouse Neural Stem Cell Nucleofector Kit (Lonza). 3-5×10^6^ NSCs were collected and resuspended in 70 μL Nucleofector solution, 15 μL supplement and 15 μL *Xist* ASO 10 μM (*XIST*-ANAND_1, cat. 339511 LG00116620-DDA: TCTTGGTTACTAACAG (8); Qiagen). Cells were nucleofected with a Lonza Nucleofector™, program A-033. Cells were then put back into culture and treated with 5-Aza or its vehicle for 3 days.

Decitabine (Selleck Chem) was resuspended in aliquots of 10 μL 10 μM in DMSO and kept in an Argon atmosphere at −80°C. During the 7-day drug test, LDN193189, GSK650394, RG2833 and decitabine were used at 0.5 μM, 2.5 μM, 5 μM and 0.5 μM respectively (8, 10). The RNA-seq analysis was performed on NSCs treated with 10 μM 5-Aza and 1.5 μM *Xist* ASO, or DMSO and 1 μM scrambled ASOs for 3 days.

### *In vitro* and *in vivo* NanoLuciferase Assays

NanoLuciferase assays were performed following manufacturer’s instructions (Nano-Glo^®^ Luciferase Assay System, Promega). Briefly, cells were collected and counted. A specific amount of cells (<0.5×10^6^ cells) were resuspended in 25 μL PBS, and added to a well of a 96-well dish (White Cliniplate, Thermo Scientific) and 25 μL of Nano-Glo^®^ Luciferase Assay Substrate + Buffer were added. Cells were lysed for 3 min and bioluminescence was then read in a VICTOR X4 Multilabel Plate Reader (1 s reading, 3 technical replicates, 10 s interval between readings).

P6 mice were anesthetized by isoflurane inhalation and injected intraperitoneally with furimazine (kindly provided by Promega). In short, solid furimazine was resuspended in 100% ethanol and kept for short periods of times at −80°C. Before injections, furimazine was resuspended in 8% glycerol, 10% ethanol, 10% hydroxypropyl-ß-cyclodextrin and 35% PEG400 in water, as in (25). Anesthetized mice were injected with 5 μg furimazine/ g mouse. Briefly after injection, mice were sacrificed and their brains extracted for imaging in an IVIS Spectrum imager (PerkinElmer). Brains were imaged in an open filter with an exposure time of 5 s (*Mecp2^+/NLucTom^*) or 5 min (*Mecp2^+/+^*). Brain radiance (p/s/cm^2^/sr) was measured from same-sized regions of interest and corrected by subtracting the background signal.

### RNA-sequencing

DNA libraries from RNA samples were prepared using the Smart-seq2 method and subsequently sequenced on an Illumina HiSeq2500 sequencer. The 50 bp single-end RNA-seq reads were processed allele-specificallyThe SNPs in the C57BL_6NJ and Cast/Ei lines were downloaded from the Sanger institute (v.5 SNP142)(26). These were used as input for SNPsplit v0.3.4 (27) to construct an N-masked reference genome based on mm10 in which all SNPs between C57BL_6NJ and Cast/Ei were masked. Reads were first mapped to a reference genome file containing the C57BL genome, Cast/Ei genome and the NanoLuciferase sequence using the default settings of hisat2 v2.2.1 (28). Reads that mapped to the NanoLuciferase sequence without mismatches were removed from the fastq files, after which the remaining reads were remapped to the N-masked reference genome. SNPsplit was then used to assign the reads to either the C57BL_6NJ or Cast/Ei bam file based on the best alignment or to a common bam file if mapping to a region without allele-specific SNPs. The allele-specific and unassigned bam files were sorted using samtools v1.10 (29). The number of mapped reads per gene were counted for both alleles separately using HTSeq v0.12.4 (--nonunique=none -m intersection-nonempty) (30) based on the gene annotation from ensembl v98. For each sample, the number of reads that mapped perfectly to the NanoLuciferase sequence was added to the *Mecp2* gene count of the Cast/Ei allele.For each condition, genes with more than 20 allele-specific reads across the triplicates were used to calculate the allelic ratio, defined as Xi/(Xi+Xa) where the inactive X (Xi) and active X (Xa) are Cast/Ei and C57BL, respectively. The difference between the allelic ratios of X-linked genes between the High and Control samples were plotted along the X chromosome using only genes with more than 20 allele-specific reads in both conditions.

We filtered the X-linked genes based on the number of reads overlapping Xa and Xi of the control samples and the High samples separately. Active genes were selected as X-linked genes with at least 6 reads overlapping Xa of the control samples, whereas inactive genes were genes with less than 6 reads overlapping the Xa^Control^. Escapee genes were selected as active genes with Xi^Control^ ≥ 5% Xa^Control^, whereas the remainder of the genes (Xi^Control^ < 5% Xa^Control^) are labelled as X-inactived genes. To find the reactivated genes, we performed a differential expression analysis using DEseq2 v1.26.0 (31), resulting in a list of genes with a significant allelic difference between High and Control. Reactivated genes were selected as X-inactivated genes that were also significantly differentially expressed (p-value < 0.05). For plotting, the counts of all genes were normalized using the variance stabilizing transformation function. For the plots with the low and medium conditions, DESeq2 was run on all four conditions where the control samples were compared against the 5-Aza samples (*i.e*. low, medium and high conditions) and counts were normalized once more. We also performed differential expression analyses between Low and Control and between Medium and Control and compared the lists of significant genes to the reactivated genes in the High samples.

The genes from the different gene classes were compared based on several characteristics. We extracted the CpG sites from the mm10 reference genome, and counted the number of CpG sites in the region ± 5kb of the TSS sites using BEDTools coverage v2.29.2 (32). For each gene, the distance from TSS to the nearest escapee and *Xist* was identified using BEDTools closest. A table containing the locations of SINE and LINE repeat elements was downloaded from UCSC and used for calculating the number of SINEs, LINEs and several SINE subtypes in the 200kb region around the TSS. Significant differences between gene classes were tested using a Mann-Whitney test with P-value < 0.05. To evaluate ChIP-seq enrichment around the TSS of the different gene groups, several publicly available ChIP-seq datasets from ESC-derived male neural progenitor cells were downloaded (CTCF, H3K4me3 and H3K27ac from GSE96107). ChIP-seq density ± 3kb around the TSS was visualized using deepTools plotProfile v3.5.0 (33).

The TAD boundaries were downloaded from Bonev et al 2017. For each TAD on the X-chromosome, the number of overlapping genes, reactivated genes and escapees were counted using bedtools intersect. TADs with significant more or less reactivated genes were selected using a Binomial test based on the ratio between the number of reactivated genes and the total number of genes for the whole X-chromosome (p-value < 0.05). For visualization, the raw NPC Hi-C read data from (14) was downloaded (GSE96107) and mapped to mm10 using bowtie2 v2.4.1 (34). For each sample, the HiC matrix was build using hicBuildMatrix from HiCExplorer v.3.6 (35) with a bin size of 10,000. The four replicates were merged into one matrix using hicSumMatrices, after which bins were merged into 50kb bins with hicMergeMatrixBins. The matrix was corrected using hicCorrectMatrix with ICE as correction method, 500 iterations and a lower and upper threshold of −3 and 6, respectively.

### MeD-seq

MeD-seq analyses were essentially carried out as previously described (12). In brief: DNA samples were digested by *LpnPI* (New England Biolabs, Ipswich, MA, USA), resulting in snippets of 32 bp around a fully-methylated recognition site that contains a CpG. These short DNA fragments were further processed using a ThruPlex DNA–seq 96D kit (cat#R400407, Rubicon Genomics Ann Arbor, MI, USA) and a Pippin system. Stem-loop adapters were blunt-end ligated to repaired input DNA and amplified to include dual indexed barcodes using a high-fidelity polymerase to generate an indexed Illumina NGS library. The amplified end product was purified on a Pippin HT system with 3% agarose gel cassettes (Sage Science, Beverly, MA, USA). Multiplexed samples were sequenced on Illumina HiSeq2500 systems for single reads of 50 bp according to the manufacturer’s instructions. Dual indexed samples were demultiplexed using bcl2fastq software (Illumina, San Diego, CA, USA). Data processing was carried out using custom scripts in Python. Raw fastq files were subjected to Illumina adaptor trimming and reads were filtered based on *LpnPI* restriction site occurrence between 13-17 bp from either 5’ or 3’ end of the read. Reads that passed the filter were mapped to mm10 using bowtie2 (28, 34). For each *LpnPI* site, the number of overlapping reads were counted and normalized for the sequencing depth. We defined the TSS region as the region ± 1kb of the TSS and generated read count scores for the TSS region of each gene. Differentially methylated TSS regions were detected using a Mann-Whitney test on the normalized read counts of the High samples and the control samples.

For each gene, the ratio between high and control was calculated by dividing the normalized number of reads overlapping the TSS region in the high samples by the those overlapping the TSS region in the control samples. Only genes with more than 10 reads overlapping the TSS region across all samples were used. The ratios between the genes of the different gene classes were compared using a violin plot showing the ratios per group. Methylation differences between the TSS region of reactivated and non-reactivated genes were explored by plotting the methylation profiles for both genes in a heatmap. For each gene, the normalized number of reads overlapping the TSS region were converted to z-scores for plotting. The genes were clustered based on the Euclidean distance and annotated as either reactivated or non-reactivated to reveal clustering differences between both groups.

For the genome browser overviews of the MeD-seq samples, the bam files were normalized using deepTools bamCoverage v3.5.0 (33) with CPM as normalization method and a bin size of 1. For each condition, the tracks of the replicates were merged using WiggleTools v1.2.3 (Zerbino et al. 2014).

### Harmony image analysis

Neurons were processed as per the IF protocol described above. Images were acquired with an Opera Phenix confocal microscope (PerkinElmer) and analyzed with a Harmony software (v4.9, PerkinElmer). Nuclei were determined with DAPI while TUJ-1-Alexa488-positive cells were selected and further analyzed for nuclear area, number of extremities per nucleus and number of roots per nucleus.

### Data access

All raw and processed high-throughput sequencing data (RNA-seq, MeD-seq) generated in this study have been submitted to the NCBI Gene Expression Omnibus (GEO) under accession number GSE166147.

